# Adipokine C1q/Tumor necrosis factor-Related Protein 3 (CTRP3) Attenuates Intestinal Inflammation via Histone Deacetylase Sirtuin 1 (SIRT1)/NF-κB Signaling

**DOI:** 10.1101/2022.05.08.491034

**Authors:** Huimin Yu, Zixin Zhang, Gangping Li, Yan Feng, Lingling Xian, Fatemeh Bakhsh, Dongqing Xu, Cheng Xu, Tyrus Vong, Bin Wu, Florin M Selaru, Fengyi Wan, G. William Wong, Mark Donowitz

**Author notes:** Address correspondence to: Huimin Yu, MD, PhD, Johns Hopkins University School of Medicine, Division of Gastroenterology and Hepatology, 720 Rutland Avenue, Ross Research Building, Room 933, Baltimore, MD 21205.

## Abstract

**BACKGROUND & AIMS:** The adipokine C1q/tumor necrosis factor-related protein 3 (CTRP3) has anti-inflammatory effects in several non-intestinal disorders. Although CTRP3 is reduced in the serum of patients with inflammatory bowel disease (IBD), its function in IBD has not been established. We aimed to elucidate the function of CTRP3 and related molecular mechanisms in intestinal inflammation using a colitis model of genetically-modified CTRP3 mice and intestinal epithelial tissue from patients with Crohn’s disease (CD), one of the two main forms of IBD.

**METHODS:** CTRP3 knockout (KO) and overexpressing transgenic (Tg) mice along with their corresponding wild-type (WT) littermates were subjected to drinking water containing dextran sulfate sodium (DSS) for 6-10 days to induce acute colitis. Mouse colitis symptoms and histological data were analyzed. CTRP3-mediated signaling was examined in the intestinal tissue of mice and patients with CD.

**RESULTS:** CTRP3 mRNA and protein were detected in murine and human intestinal epithelial cells, as well as in murine intestinal smooth muscle cells and mesenteric fat. In DSS-induced acute colitis models, CTRP3 KO mice developed more severe colitis than their WT littermates, while CTRP3 overexpressing Tg mice developed less severe colitis than their WT littermates. In both water- and DSS-treated CTRP3 KO mice, reduced CTRP3 levels correlated with decreased levels of Sirtuin 1 (SIRT1), a histone deacetylase, increased levels of phosphorylated nuclear factor kappa B (NF-κB) subunit p65, resulting in increased expression of pro-inflammatory cytokines tumor necrosis factor-α (TNF-α) and interleukin 6 (IL-6). The results from CTRP3 Tg mice mirrored those from CTRP3 KO mice in most respects. This CTRP3/SIRT1/NF-κB relationship was also observed in the intestinal epithelial tissue of patients with active and inactive CD.

**CONCLUSIONS:** CTRP3 expression levels correlate negatively with intestinal inflammation in mouse colitis models and CD patients. CTRP3 attenuates intestinal inflammation via SIRT1/NF-κB signaling to suppress pro-inflammatory cytokines in mouse colitis models and patients with IBD. The manipulation of CTRP3 signaling, including through the use of SIRT1 agonists, may offer translational potential in the treatment of IBD.

**WHAT YOU NEED TO KNOW:** *BACKGROUND AND CONTEXT:* Adipokine C1q/tumor necrosis factor-related protein 3 (CTRP3) is a novel adipokine with known non-intestinal anti-inflammatory effects. CTRP3 is reduced in the serum of patients with inflammatory bowel disease (IBD). However, little is known about whether and how CTRP3 influences intestinal inflammation in IBD.

*NEW FINDINGS:* CTRP3 mRNA and protein were detected in murine and human intestinal epithelial cells, as well as in murine intestinal smooth muscle cells and mesenteric fat. CTRP3 deletion was associated with more severe acute dextran sulfate sodium (DSS)-induced colitis, while CTRP3 overexpression was associated with less severe colitis. In both mice and humans, reduced CTRP3 levels correlated with reduced levels of the histone deacetylase Sirtuin 1 (SIRT1), resulting in the up-regulation of phosphorylated nuclear factor-kappa B (NF-κB) p65 and pro-inflammatory cytokine synthesis.

*LIMITATIONS:* This study was performed using genetically modified mice and human tissue samples. An acute DSS-induced colitis model was used; additional mouse colitis models designed to mimic other aspects of IBD will be examined in future studies. The specific source of the secreted CTRP3 protein which influences intestinal inflammation is yet to be identified. The use of recombinant CTRP3 protein supplementation and SIRT1 agonists to mitigate intestinal inflammation also requires further study.

*IMPACT:* CTRP3 is a novel anti-inflammatory adipokine that attenuates intestinal inflammation in colitis mouse models and intestinal epithelial tissue of patients with IBD. CTRP3 attenuates intestinal inflammation by activating SIRT1, which suppresses the pro-inflammatory transcriptional activity of phosphorylated NF-κB p65. CTRP3 and SIRT1 agonists have potential as novel IBD drug targets.

## Introduction

Inflammatory bowel disease (IBD), including Crohn’s disease (CD) and ulcerative colitis (UC), is a chronic, debilitating inflammatory disorder of the gastrointestinal tract that affects 3 million people in the US (1.3% of adults).^1–3^ Despite advances brought on by recently developed biologics, including TNF-α inhibitors, nearly one-third of patients with IBD have failed to respond to existing therapies.^4–7^ There is a pressing need to develop evidence-based, novel strategies to control refractory inflammation in IBD.

Mesenteric fat and its secreted mediators represent a promising area in IBD research.^8–12^ Since the earliest descriptions of CD, the presence of hyperplastic mesenteric fat that characteristically wraps itself around the inflamed intestine, typically the terminal ileum, has been noted as an indicator of severe disease.^13–15^ This so-called “creeping fat” correlates with intestinal transmural inflammation, muscular hypertrophy, and fibrosis/stricture formation.^16–20^ Despite its longstanding association with disease progression, the precise function of mesenteric fat remains obscure. Mesenteric fat is likely the principal source of adipokines and cytokines responsible for inflammatory processes associated with IBD.^16–18^ And yet it appears to serve as a barrier to inflammation, controlling immune responses to translocated gut bacteria.^19^ The complex interplay between mesenteric fat and intestinal inflammation in IBD has, understandably, attracted growing interest.^21–25^

Among the mediators secreted by mesenteric (and creeping) fat, <u>C</u>1q/<u>T</u>umor necrosis factor-<u>R</u>elated <u>P</u>rotein 3 (CTRP3) is of particular interest.^26–29^ CTRP3 was initially characterized in 2004 and was noted for containing a C1q globular domain similar to adiponectin and tumor necrosis factor (TNF), both of which are known to be involved in intestinal inflammation.^30–36^ Since then, CTRP3 has been found to correlate negatively with several pro-inflammatory cytokines, including TNF-α, interleukin 6 (IL-6), IL-8, and C-reactive protein (CRP).^33, 34, 37–41^ CTRP3 has been shown to attenuate inflammation in several non-intestinal conditions, including obesity,^35,41–44^ diabetes,^35,37,39,45^ rheumatoid arthritis,^46^ acute pancreatitis,^47^ and cardiac inflammation.^48,49^ Notably, the serum levels of CTRP3 are reduced in IBD patients.^50^ The function of CTRP3 in IBD, however, has not previously been established.

CTRP3 appears to signal through different pathways in different tissue/cell types.^27^ Two possible CTRP3 receptors have been suggested, but none has been confirmed.^51^ In settings of pancreatitis and cardiac inflammation, CTRP3 was found to exert its anti-inflammatory effect by inhibiting nuclear factor kappa B (NF-κB) signaling via Sirtuin 1 (SIRT1), a histone deacetylase, thereby down-regulating pro-inflammatory cytokine production.^47,49^ SIRT1/NF-κB signaling is known to influence intestinal inflammation in IBD^52–54^ and SIRT1 levels are known to be reduced in the colonic tissue of IBD patients and mice with dextran sulfate sodium (DSS)-induced colitis.^55–57^

We hypothesized that CTRP3 attenuates intestinal inflammation in IBD through SIRT1/NF-κB signaling. We demonstrated that CTRP3 expression levels correlate negatively with intestinal inflammation in acute DSS-induced mouse colitis models and CD patients. We also showed that, in both mice and humans, CTRP3 attenuates intestinal inflammation through SIRT1, which suppresses NF-κB transcriptional activity, resulting in reduced production of pro-inflammatory cytokines. Our findings raise the possibility that the administration of CTRP3 protein and/or SIRT1 agonists may represent a novel treatment approach to IBD. More broadly, our findings add to the growing body of evidence that adipokines are important to IBD pathogenesis, and may assist in the development of new, adipokine-based treatments for IBD.

## Materials and Methods

### CTRP3 knockout (KO) and overexpressing transgenic (Tg) mice

All animal protocols were approved by the Institutional Animal Care and Use Committee of the Johns Hopkins University School of Medicine. CTRP3 KO (-/-) and overexpressing Tg mouse strains were previously generated.^34,36^ CTRP3 KO mice and wild-type (WT) littermates were generated by intercrossing CTRP3 heterozygous (+/-) mice. Littermates were used throughout the study. Upon termination of the study, animals were fasted overnight and euthanized; blood was collected for serum, and tissues were collected and snap-frozen in liquid nitrogen. The above samples were kept at −80 °C until analysis. All mice were co-housed to control for microbiome differences.

### Human samples

Human intestinal epithelial tissue and mesenteric fat were obtained from CD and control patients undergoing endoscopic or surgical procedures through two research protocols approved by the Johns Hopkins Institutional Review Boards. Tissue was de-identified, but age, gender, and ethnicity information were noted.

### Acute DSS colitis mouse models

Age- and sex-matched CTRP3 KO and Tg mice, along with their WT littermates, were subjected to different concentrations of DSS (1.2%, 1.5%, 2.0%, and 2.5%) in drinking water for 6-10 consecutive days. Mouse body weight, stool consistency, and rectal bleeding were recorded and scored daily as shown in Supplementary Table 1.^58^ Surviving analysis was based on clinical data collected over 10 days of DSS treatment. Upon termination of the study, blood was collected for serum, and mice were dissected for tissues of interest (mesenteric fat, terminal ileum, and colon). Tissue was fixed in 4% paraformaldehyde (PFA) overnight followed by paraffin embedding. 5-8 μm thick paraffin sections were stained with Hematoxylin and Eosin (H&E) and examined in a blinded manner by two researchers (YF and HY). Tissue damage and inflammatory cell infiltrates were scored as shown in Supplementary Table 2.^59^

### Single-molecule RNA fluorescence in situ hybridization (smFISH) for RNA detection in mouse intestine

smFISH employs a probe library consisting of short DNA oligonucleotides labeled with a fluorescent dye that hybridize to a target RNA in fixed cells, allowing RNA quantification and localization at the single-cell level and with single-molecule resolution.^60^ All buffers were prepared following RNase precautions and using RNase-free water. Briefly, intestinal tissue was fixed in 4% PFA overnight and washed three times with phosphate-buffered saline (PBS). The tissue was then embedded in the Optimal Cutting Temperature (OCT) compound, cut into 5-10 μm frozen sections, and mounted onto collagen-coated coverslips. The sections were washed with 1X PBS and incubated in 70% alcohol overnight at 4 °C. After removing 70% alcohol and submerging them in 15% formamide and 2X SSC, they were incubated in pre-hybridization buffer for 20 minutes. Then, sections were incubated in 50 μl of hybridization solution containing 80 nM DNA probe libraries for mouse CTRP3 (a mixture of 55 cy3-labeled oligonucleotides) and mouse glyceraldehyde 3-phosphate dehydrogenase (GAPDH, a mixture of 33 cy5-labeled oligonucleotides) in a humidified chamber overnight at 37 °C. After that, sections were rinsed with pre-warmed hybridization buffer and incubated for 20 minutes at 37 °C. Then, sections were incubated again in the hybridization buffer for 20 minutes at room temperature. Finally, sections were washed with 1X PBS two times and mounted on slides with ProLong Diamond Antifade Mountant (Thermo Fisher Scientific). Images were taken and analyzed using an automated inverted Nikon Ti-2 wide-field microscope equipped with x 60 1.4NA oil immersion objective lens (Nikon), Spectra X LED light engine (Lumencor), and ORCA-Flash 4.0 V2 sCMOS camera (Hamamatsu).

### Immunostaining of mouse and human intestine

Mouse intestinal tissue with attached mesenteric fat or human terminal ileal epithelial tissue was fixed overnight at 4% PFA and then embedded in the OCT compound. Tissue staining was performed according to standard immunofluorescence protocols with the following primary antibodies: mouse CTRP3 (generated and provided by GWW, rabbit anti-mouse), human CTRP3 (Invitrogen, PA5-115061, rabbit anti-human), α-SMA-Cy3 (Sigma, A2547). For detection, the following fluorescent-labeled secondary antibodies were used: Alexa Fluor^®^ 647 phalloidin (Molecular Probes, A22287); Alexa 488 donkey anti-rabbit IgG (Molecular Probes, A11008); Alexa 594 goat anti-rabbit IgG (Molecular Probes, A11012). The tissue was counterstained using 4’,6-diamidino-2-phenylindole (Sigma, D8417). Images were taken and analyzed using a FISHscope at the Johns Hopkins Ross Fluorescence Imaging Center.

### Reverse transcription-quantitative polymerase chain reaction (RT-qPCR)

mRNA gene expression of CTRP3, SIRT1, TNF-α, and IL-6 were measured by RT-qPCR. Total RNA of mouse and human intestinal tissue was isolated using the RNeasy Mini Kit (Qiagen) and total RNA of mouse mesenteric fat was isolated using the RNeasy Lipid Tissue Mini Kit (Qiagen). The isolated RNA was transcribed to complementary DNA via Superscript IV VILO Master Mix (Invitrogen). Quantitative PCR was performed with the PowerTrack SYBR Green Master Mix (Thermo Fisher Scientific). All qPCR primers used are listed in Supplementary Table 3. Data were analyzed by the comparative cycle threshold (Ct) method as means of relative quantitation of gene expression, normalized to an endogenous reference control gene (mouse beta-2-microglobulin or human beta-actin) and relative to a calibrator (normalized Ct value obtained from WT mice or control patients), and expressed as 2^-ΔΔCT^.

### Western blot analysis

Mouse or human intestinal tissue was lysed in 100 μl ice-cold RIPA buffer (Thermo Fisher Scientific) with protease inhibitors (Thermo Fisher Scientific). Homogenates were incubated in a cold room for 2 hours and then centrifuged at 16000 *g* for 20 minutes at 4 °C. 10-15 μg proteins of the whole-cell extracts were run on 10% sodium dodecyl sulfate (SDS) polyacrylamide gel for 1 hour at 150V. Proteins were then transferred to nitrocellulose membranes for 1.5 hours at 100V. The membranes were blocked with PBS supplemented with 5% dry milk for 1 hour at room temperature, probed with the following primary antibodies: β-actin (Sigma, A1978,), mCTRP3 (provided by GWW), hCTRP3 (Invitrogen, PA5-115061), SIRT1 (Cell Signaling, #2028 and #8469), NF-κB p65 (Santa Cruz, sc-372), p-NF-κB p65 (Cell Signaling, #3033) at 4 °C overnight, and then washed in TBST for 30 minutes. The membranes were probed with corresponding secondary antibodies at 1:10000 dilution for 1 hour at room temperature and the bands were visualized by a Li-Cor Odyssey Fc imaging system and Image Studio software. Signals were quantified using ImageJ (National Institutes of Health, Bethesda, MD) and normalized to β-actin.

### Statistical analysis

All data presented are representative of three or more independent experiments and shown as mean ± SEM unless stated otherwise. The difference between the two groups was assessed by Student’s t-test. The difference among three or more groups was compared using two-way ANOVA with Tukey’s multiple comparisons test. Log-rank (Mantel-Cox) testing was used for survival analysis. Statistical analyses were performed using GraphPad Prism software. A statistically significant difference was considered when \**P* < .05, \*\**P* < .01, \*\*\**P* < .001, and \*\*\*\**P* < .0001.

Additional experimental details are available in the Supplementary Material.

## Results

### CTRP3 Localizes to Murine Intestinal Epithelial Cells, Intestinal Smooth Muscle Cells, and Mesenteric Fat

Immunofluorescence staining of colonic tissue from wild-type (WT) mice detected CTRP3 protein in intestinal epithelial cells, intestinal smooth muscle cells, and mesenteric adipocytes (Figure 1A). Colonic tissue from CTRP3 KO mice was used as a negative control (Supplementary Figure 1A). Single-molecule RNA fluorescence *in situ* hybridization (smFISH) of WT colonic tissue confirmed CTRP3 mRNA expression in all three cell types (Figure 1B, Supplementary Figure 1B). Notably, CTRP3 mRNA and protein expression levels were higher in the colon than in mesenteric fat (Figure 1C and D).

**Figure 1.**
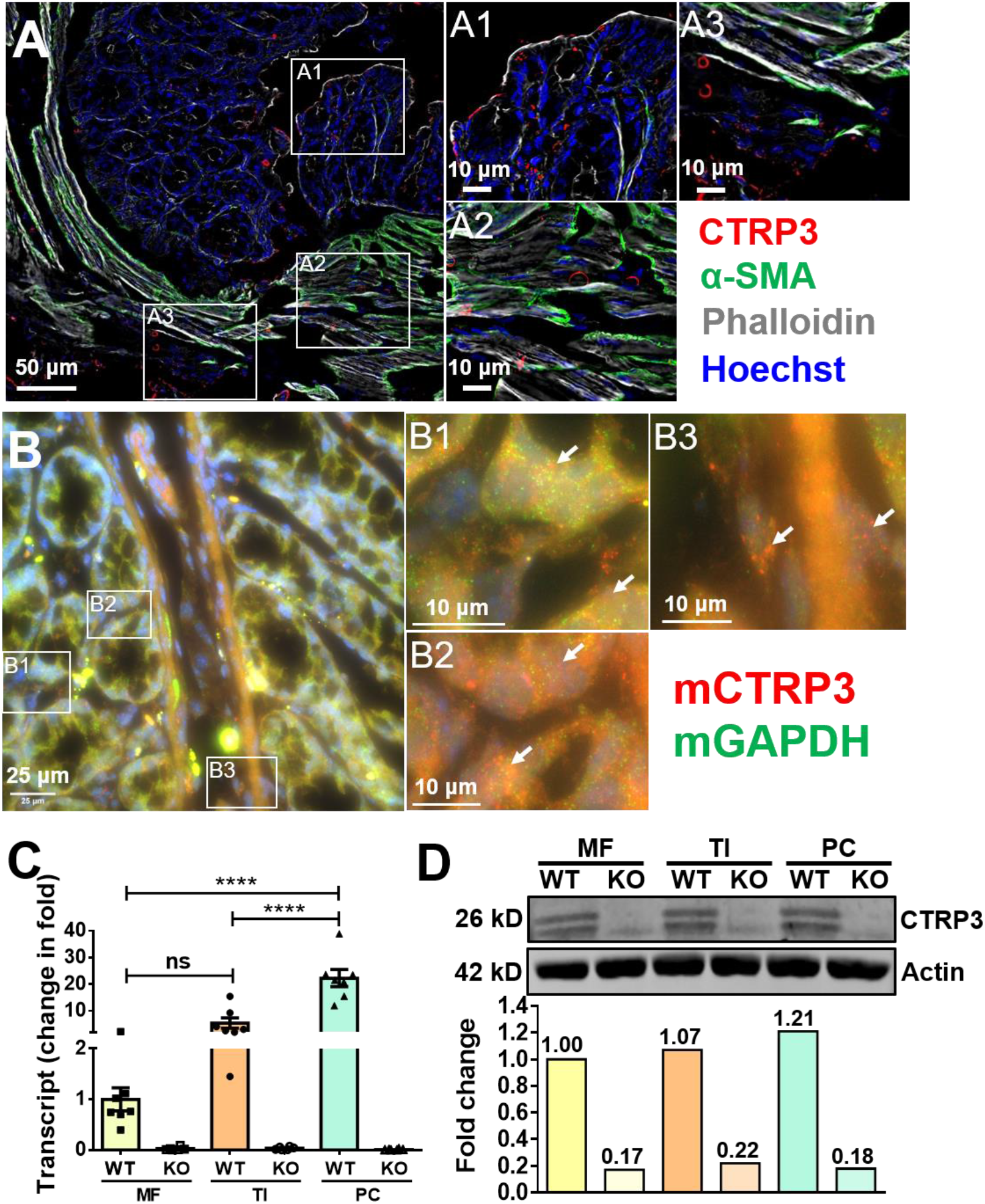
CTRP3 is expressed in mouse intestinal epithelial cells, intestinal smooth muscle cells, and mesenteric fat. **(A)** Immunofluorescence staining of WT colonic tissue showed that CTRP3 protein was expressed in intestinal epithelial cells (A1), intestinal smooth muscle cells (A2), and mesenteric fat (A3) attached to the colon. α-SMA (smooth muscle actin), Phalloidin (F-actin), and Hoechst (nuclei). Experiments were repeated three times. **(B)** Single-molecule RNA fluorescence *in situ* hybridization (smFISH) of WT colonic tissue showed that CTRP3 mRNA (red dots) was expressed in intestinal epithelial cells (B1 and B3) and intestinal smooth muscle cells (B2). White arrows indicate cells expressing CTRP3 mRNA (red dots). GAPDH mRNA (green dots, a positive control for smFISH). **(C)** RT-qPCR analysis of CTRP3 mRNA expression levels in the mesenteric fat (MF), terminal ileum (TI), and proximal colon (PC) of CTRP3 KO and WT mice. Results are shown as means ± SEM; n = 6-7 mice/group. Results are shown as means ± SEM; n = 6mice/group. \**P* < .05; \*\**P* < .01; \*\*\**P* < .001; and **** *P* < .0001 (two-way ANOVA). **(D)** A representative immunoblot of CTRP3 protein expression levels in the mesenteric fat (MF), terminal ileum (TI), and proximal colon (PC) of CTRP3 KO and WT mice. Protein bands were quantified using ImageJ and normalized to β-actin.

### Deletion of CTRP3 Aggravates Intestinal Inflammation and Tissue Injury in Acute DSS-induced Mouse Colitis

To determine whether CTRP3 plays a role in intestinal inflammation *in vivo,* we established a DSS-induced acute colitis model of CTRP3 KO mice and their corresponding WT littermates. CTRP3 KO mice were viable, fertile, and developed normally with no gross abnormal phenotype.^36^ The CTRP3 KO mice were found at the expected Mendelian frequency at birth. After 6 days of DSS treatment, the mesenteric fat from WT mice displayed adipocyte hyperplasia, with smaller but more abundant adipocytes, and more inflammatory cell infiltration, compared with that from water-treated controls (Supplementary Figure 2). These histologic changes are consistent with what has been reported in creeping fat^10,40^ and confirmed in our own human studies (Figure 6B-E).

Four different concentrations of DSS (1.2%, 1.5%, 2.0%, and 2.5%) in drinking water were tested for 6-10 consecutive days, with 1.2% DSS treatment resulting in the most distinguishable colitis phenotypes between CTRP3 KO and WT mice. 1.2% DSS was therefore used in subsequent experiments (Figure 2A). After 7 days of 1.2%DSS treatment, both CTRP3 KO and WT mice developed obvious colitis, while the corresponding water-treated control groups showed no clinical or histological signs of colitis (Figure 2B-D, F-G). Compared to the DSS-treated WT mice, DSS-treated CTRP3 KO mice developed significantly more severe colitis as evidenced by more pronounced weight loss (Figure 2B), higher colitis disease scores (Figure 2C), and a lower survival rate (Figure 2D). Histologic analysis of the proximal and distal colon revealed more tissue damage and inflammatory cell infiltrates in DSS-treated CTRP3 KO mice than in DSS-treated WT mice (Figure 2F-G, Supplementary Figure 3A). In addition, the colons of DSS-treated KO mice were also significantly shorter than those of DSS-treated WT mice (Figure 2E; 5.45 ± 0.16 cm vs. 6.56 ± 0.28 cm, *P* < .05).

**Figure 2.**
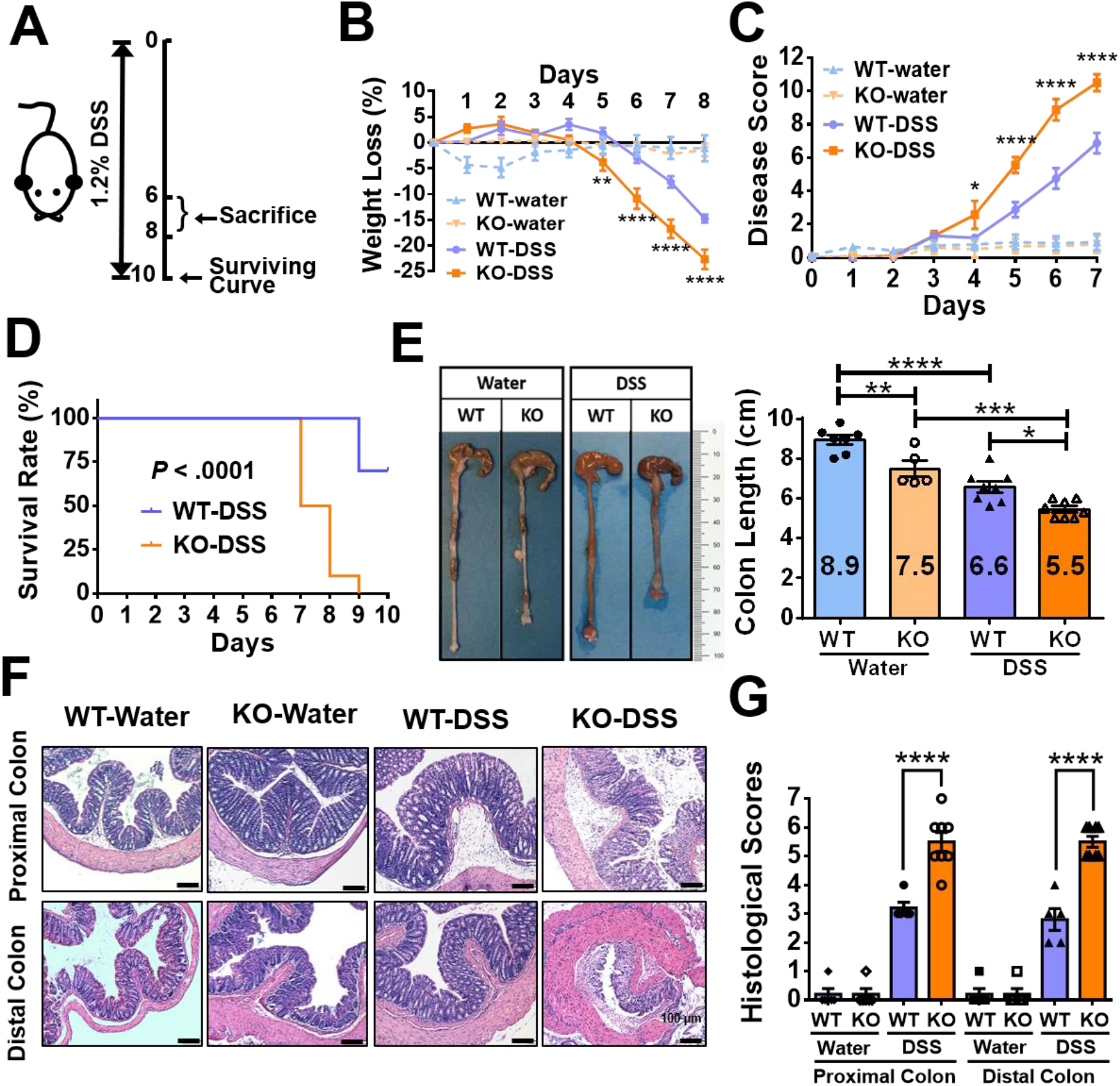
CTRP3 KO mice develop more severe acute colitis than WT littermates in acute DSS-induced colitis models. **(A)** Experimental scheme for CTRP3 KO DSS-induced acute colitis. 1.2% DSS was administered in drinking water for 6-10 days to induce acute colitis. **(B-C)** DSS-treated CTRP3 KO mice had more pronounced weight loss and developed more severe colitis relative to DSS-treated WT littermates. Water-treated CTRP3 KO and WT mice showed no difference in weight loss and clinical colitis phenotypes. Statistical comparison was between DSS-treated CTRP3 KO and WT mice. **(D)** DSS-treated CTRP3 KO mice displayed a lower survival rate relative to DSS-treated WT littermates. Log-rank Mantel-Cox test was used for survival analysis. **(E)** The colons of CTRP3 KO mice were significantly shorter than those of WT littermates in both water- and DSS-treated groups. **(F-G)** H&E staining of proximal and distal colon revealed more tissue damage and inflammatory cell infiltrates in DSS-treated CTRP3 KO mice than in DSS-treated WT mice. The histological scores were not different between water-treated CTRP3 KO and WT mice. Scale bar, 100 *μ*m. Results are shown as means ± SEM; n = 5-10 mice/group. \**P* < .05; \*\**P* < .01; \*\*\**P* < .001; **** *P* < .0001 (twoway ANOVA except for the survival curve). The above data are from male CTRP3 KO and WT mice; female data were similar (Supplementary Figure 4).

The inflammatory effects of CTRP3 deletion were less evident under basal conditions (water treatment) than when colitis was induced (DSS treatment), but nonetheless were detected. Water-treated CTRP3 KO and WT mice showed no difference in clinical colitis phenotypes (Figure 2B-D), and histologic analysis likewise revealed no meaningful differences (Figure 2F-G, Supplementary Figure 3A). However, the colons of water-treated CTRP3 KO mice were significantly shorter than those of WT littermates (Figure 2E; 7.48 ± 0.40 cm vs. 8.93 ± 0.24 cm, *P* < .01).

### Overexpression of CTRP3 Attenuates Intestinal Inflammation and Tissue Injury in Acute DSS-induced Mouse Colitis

To further confirm the anti-inflammatory function of CTRP3 in intestinal inflammation, we developed a DSS-induced acute colitis model of CTRP3 overexpressing Tg mice and their corresponding WT littermates. Like the CTRP3 KO mice, the CTRP3 Tg mice were viable, fertile, and did not display any gross phenotype under baseline conditions.^34^ Again, several concentrations of DSS in drinking water were tested for 6-10 days, here with 2% DSS treatment best distinguishing the colitis phenotypes between CTRP3 Tg and WT mice (Figure 3A). After 7 days of 2% DSS treatment, CTRP3 Tg mice developed significantly less severe colitis than their DSS-treated WT mice, experiencing less pronounced weight loss (Figure 3B), lower colitis disease scores (Figure 3C), and a higher survival rate (Figure 3D). Histological analysis of the proximal and distal colon showed that DSS-treated CTRP3 Tg mice exhibited significantly less tissue damage and fewer inflammatory cell infiltrates than their WT littermates (Figure 3F-G, Supplementary Figure 3B). Further, post-mortem examination revealed that the colons of DSS-treated CTRP3 Tg mice were significantly longer than those of their DSS-treated WT littermates (Figure 3E; 8.42 ± 0.08 cm vs. 7.71 ± 1.05 cm, *P* < .001).

**Figure 3.**
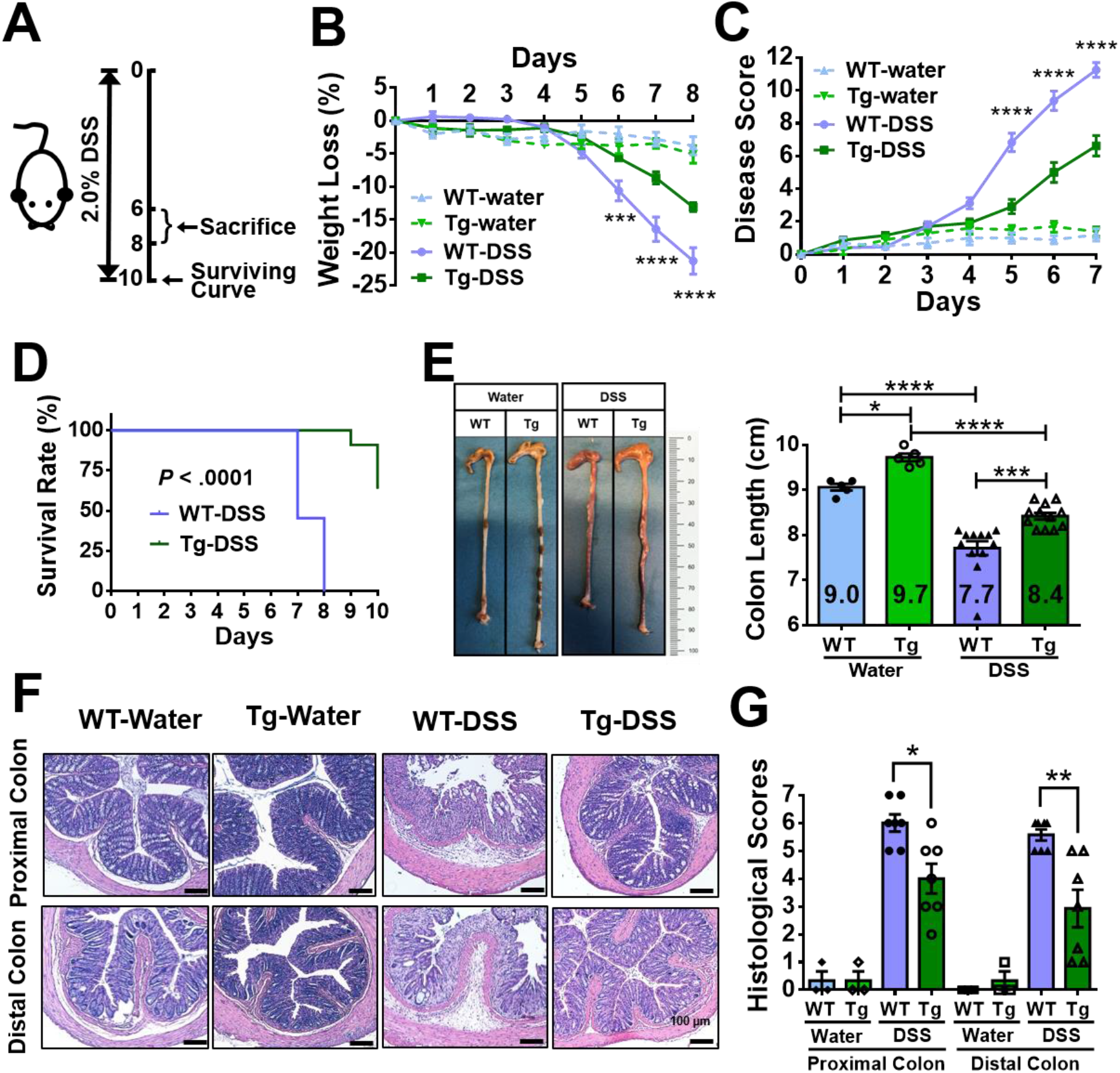
CTRP3 overexpressing Tg mice develop less severe acute colitis than WT littermates in acute DSS-induced colitis models. **(A)** Experimental scheme for CTRP3 Tg DSS-induced acute colitis. 2% DSS was administered in drinking water for 6-10 days to induce acute colitis. **(B-C)** DSS-treated CTRP3 Tg mice had less pronounced weight loss and developed less severe colitis relative to DSS-treated WT littermates. Water-treated CTRP3 Tg and WT mice exhibited similar weight loss and colitis disease scores. Statistical comparison was between DSS-treated CTRP3 Tg and WT mice. **(D)** DSS-treated CTRP3 Tg mice displayed a higher survival rate relative to DSS-treated WT littermates. Log-rank Mantel-Cox test was used for survival analysis. **(E)** The colons of CTRP3 Tg mice were significantly longer than those of WT littermates in both water- and DSS-treated groups. **(F-G)** H&E staining of proximal and distal colon revealed less tissue damage and inflammatory cell infiltrates in DSS-treated CTRP3 Tg mice than in DSS-treated WT mice. The histological scores were not different between water-treated CTRP3 Tg and WT mice. Scale bar, 100 *μ*m. Results are shown as means ± SEM; n = 5 −12 mice/group. \**P* < .05; \*\**P* < .01; \*\*\**P* < .001; **** *P* < .0001 (two-way ANOVA except for the survival curve). The above data are from male CTRP3 Tg and WT mice; female data were similar (Supplementary Figure 5).

The effects of CTRP3 overexpression under basal conditions (water treatment) mirrored the effects of CTRP3 deletion under basal conditions. That is, water-treated CTRP3 Tg and WT mice showed no differences in clinical colitis phenotypes (Figure 3B-D), nor did histologic analysis reveal meaningful differences (Figure 3F-G, Supplementary Figure 3B), but the colons of water-treated CTRP3 Tg mice were significantly longer than those of WT littermates (Figure 3E; 9.72 ± 0.09 cm vs. 9.06 ± 0.07 cm, *P* < .05). Thus, CTRP3 expression levels were positively correlated with mouse colon length in both basal (water treatment) and colitis (DSS treatment) conditions in both CTPR KO and Tg mice, compared with their corresponding WT littermates.

### Deletion of CTRP3 Reduces Intestinal SIRT1 and Up-regulates Phosphorylated NF-κB p65 and Inflammatory Cytokines in Acute DSS-induced Mouse Colitis

Next, we investigated whether CTRP3 attenuates intestinal inflammation through SIRT1/NF-κB signaling in CTRP3 KO mice and WT mice. Specifically, we tested our hypothesis that CTRP3 attenuates intestinal inflammation through SIRT1, which suppresses the pro-inflammatory transcriptional activity of phosphorylated NF-κB p65. We examined the colonic expression levels of CTRP3, SIRT1, phosphorylated NF-κB p65, and pro-inflammatory cytokines TNF-α, and IL-6 in both CTRP3 KO and WT mice after DSS and water treatment.

Interestingly, 7 days of DSS treatment correlated with a significant reduction in CTRP3 levels in WT mouse colons (Figure 4A, E-F). As expected, colonic CTRP3 levels were undetectable in both water- and DSS-treated CTRP3 KO mice (Figure 4A, E-F). Compared with WT mice, SIRT1 levels were lower in CTRP3 KO mice in both the water- and DSS-treated groups (Figure 4B, E-F). In all four groups, NF-κB p65 levels were similar in the colonic whole-cell extracts of all four groups; however, phosphorylated NF-κB p65 levels were significantly up-regulated in CTRP3 KO mice relative to WT mice, and higher in DSS-treated CTRP3 KO mice than in water-treated CTRP3 KO mice (Figure 4E-F). As expected, colonic mRNA levels of TNF-α and IL-6 were up-regulated after DSS treatment in both WT and CTRP3 KO mice (Figure 4C-D). Compared to DSS-treated WT mice, TNF-α and IL-6 were significantly up-regulated in the DSS-treated CTRP3 KO mice (Figure 4C-D). Notably, in the water-treated groups, TNF-α and IL-6 were up-regulated in both the colons and terminal ilea of CTRP3 KO mice compared with WT mice (Figure 4C-D, two-tailed Student’s t-tests, *P* = 0.0001 for TNF-α and *P* = 0.0014 for IL-6; Supplemental Figure 6).

**Figure 4.**
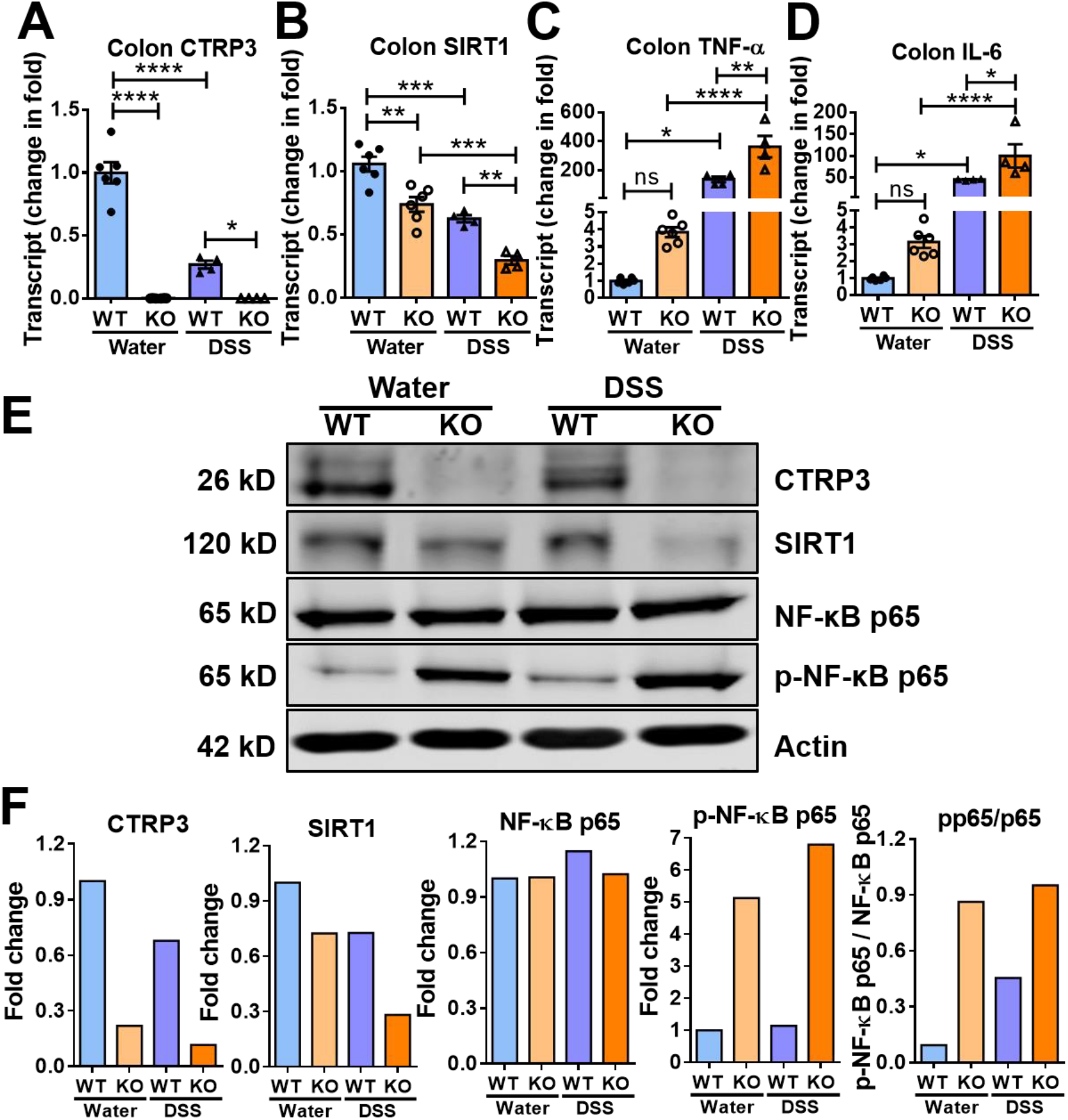
CTRP3 KO mice exhibit decreased intestinal SIRT1 and increased phosphorylated NF-κB p65 transcriptional activity in acute DSS-induced mouse colitis. Colons were used in all experiments. **(A)** CTRP3 mRNA levels were reduced in the colons of WT mice after 7 days of 1.2% DSS administration. **(B)** SIRT1 mRNA levels were reduced in the colons of water-treated CTRP3 KO mice relative to those of WT mice. DSS treatment further reduced the SIRT 1 mRNA levels in the colons of both CTRP3 KO mice and WT mice, more severely in CTRP3 KO mice than WT mice. (**C-D)** TNF-α and IL-6 mRNA levels were up-regulated in CTRP3 KO mice compared to WT mice (DSS- and water-treated). **(E-F)** CTRP3 protein levels were reduced in the colons of DSS-treated WT mice relative to those of water-treated WT mice. SIRT1 protein levels were reduced in the colons of CTRP3 KO mice than in those of WT mice, more severely in the DSS-treated mice than in the water-treated KO mice. NF-κB p65 from colonic whole-cell extracts had similar expression levels across all four groups; however, phosphorylated NF-kB p65 was up-regulated in the colons of CTRP3 KO mice, more severely in DSS-treated than water-treated KO mice. Protein bands were quantified using ImageJ and normalized to β-actin (F). Results are shown as means ± SEM; n = 6 mice/group. \**P* < .05; \*\**P* < .01; \*\*\**P* < .001; **** *P* < .0001; ns, not significant (two-way ANOVA).

In sum, we showed that in both acute colitis (DSS treatment) and basal conditions (water treatment), CTRP3 KO mice expressed reduced levels of SIRT1 and up-regulated phosphorylated NF-κB p65 compared with WT mice, resulting in increased intestinal TNF-α and IL-6. These data suggest that CTRP3 exerts an anti-inflammatory effect in acute colitis models through SIRT1, which suppresses pro-inflammatory NF-κB transcriptional activity.

### Overexpression of CTRP3 Increases Intestinal SIRT1 and Down-regulates Phosphorylated NF-κB p65 and Inflammatory Cytokines in Acute DSS-induced Mouse Colitis

To further confirm that CTRP3 attenuates intestinal inflammation through SIRT1/NF-κB signaling, we performed similar experiments in water- and DSS-treated CTRP3 overexpressing Tg and WT mice. Once again, 7 days of DSS treatment correlated with a significant reduction in CTRP3 levels in WT mouse colons (Figure 5A, E-F, *P* = 0.0008 for CTRP3 mRNA, two-tailed Student’s t-tests). As anticipated, in water- and DSS-treated CTRP3 Tg mice, colonic CTRP3 levels were significantly higher relative to WT mice (Figure 5A, E-F). SIRT1 levels were likewise higher in water-treated CTRP3 Tg mice than in water-treated WT mice, while DSS-treatment reduced SIRT1 to similar levels in both CTRP3 Tg and WT mice (Figure 5B, E-F). NF-κB p65 levels were similar in the colonic whole-cell extracts of all four groups (Figure 5E-F). Consistent with our findings in our CTRP3 KO mice experiments, however, phosphorylated NF-κB p65 levels were down-regulated in DSS-treated CTRP3 Tg mice compared with DSS-treated WT mice (and higher in the DSS group than in the water group). As expected, colonic mRNA levels of TNF-α and IL-6 were up-regulated after DSS treatment in both WT and CTRP3 Tg mice (Figure 5C-D). Compared to DSS-treated WT mice, TNF-α and IL-6 were significantly down-regulated in the DSS-treated CTRP3 Tg mice (Figure 5C-D). In the water-treated groups, levels of TNF-α were lower in CTRP3 Tg mice compared with WT mice, while the difference in IL-6 levels was not statistically significant (Figure 5C-D, two-tailed Student’s t-tests, *P* = 0.0178 for TNF-α and *P* = 0.3702 for IL-6, Student’s t-test).

**Figure 5.**
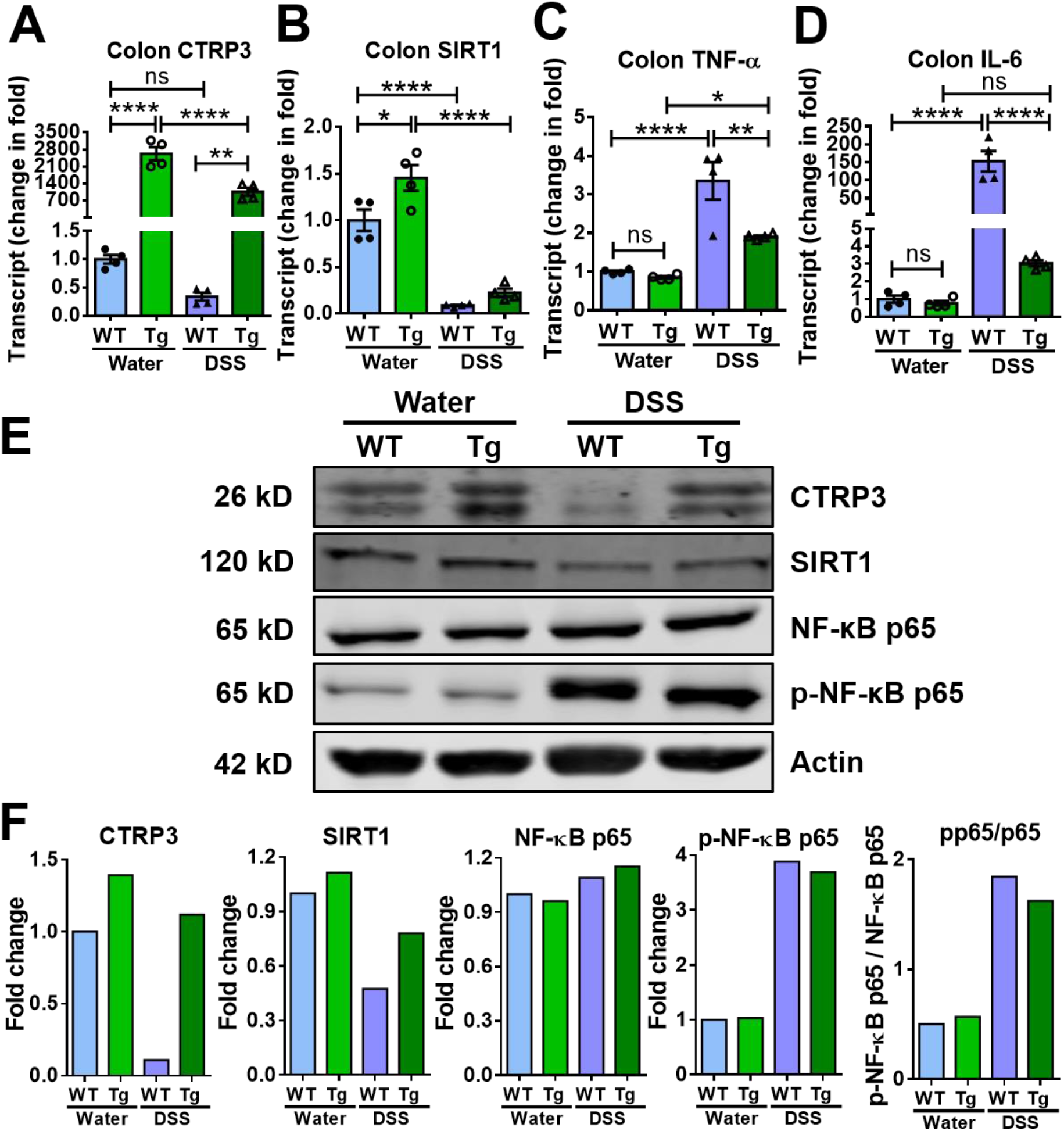
CTRP3 Tg mice exhibit increased intestinal SIRT1 and decreased phosphorylated NF-κB p65 transcriptional activity in acute DSS-induced mouse colitis.Colons were used in all experiments. **(A)** CTRP3 mRNA levels were significantly reduced in both CTRP3 Tg and WT mice after 7 days of 2% DSS administration. **(B)** SIRT1 mRNA levels were higher in CTRP3 Tg mice than in WT mice (water-treated). DSS treatment reduced SIRT1 mRNA in both CTRP3 Tg and WT mice. (**C-D)** TNF-α and IL-6 mRNA levels were down-regulated in CTRP3 Tg mice compared to WT mice (DSS-treated). **(E-F)** SIRT1 protein levels were higher in CTRP3 Tg relative to WT mice (both water-treated and DSS-treated). NF-κB p65 from the colonic whole-cell extracts had similar expression levels across all four groups; however, phosphorylated NF-κB p65 was significantly higher in the DSS-treated mice than in the water-treated mice. The ratio of phosphorylated NF-κB p65/ NF-κB p65 was reduced in CTRP3 Tg mice relative to WT mice (DSS-treated). Protein bands were quantified using ImageJ and normalized to β-actin (F). Results are shown as means ± SEM; n = 4 mice/group. \**P* < .05; \*\**P* < .01; \*\*\**P* < .001; **** *P* < .0001; ns, not significant (two-way ANOVA).

These CTRP3 overexpression findings mirror our CTRP3 deletion findings in most respects. In both acute colitis models and under basal conditions, CTRP3 overexpression correlated with increased SIRT1 levels, down-regulated phosphorylated NF-κB p65, and decreased expression of inflammatory cytokines in the colons. Taken together, these CTRP3 KO and Tg data suggest that CTRP3 attenuates intestinal inflammation in mouse colitis models through SIRT1/NF-κB signaling.

### Reduced CTRP3 in CD Patients Down-regulates SIRT1 and Up-regulates NF-κB p65 Transcriptional Activity

We next investigated the anti-inflammatory function of CTRP3 in CD patients. Three types of terminal ileal epithelial tissue were obtained from the patients who underwent endoscopic or surgical procedures: (1) disease-affected mucosa with discernable ulcers/erosions from CD patients (active; n = 8); (2) normal-appearing mucosa from the same CD patients (inactive; n = 8); and (3) normal-appearing mucosa from control patients (control, normal histology; n = 9). The terminal ileum was chosen because it is the most commonly affected intestinal segment to develop creeping fat in CD (Figure 6A).^23^ Consistent with our observations in the murine experiments (Supplementary Figure 2), and as previously reported in human studies,^10,15^ creeping fat from CD patients contained smaller but more abundant adipocytes than the mesenteric fat from control patients, and was distinguished by marked inflammatory cell infiltrates and collagen depositions (Figure 6B-E). Immunofluorescence staining showed that CTRP3 protein levels in the terminal ileal mucosa of active CD patients were substantially reduced, compared to control patients (Figure 6F-H). RT-qPCR analysis of terminal ileal mucosa showed that mRNA levels of both CTRP3 and SIRT1 were reduced in CD patients, with the difference between the inactive and active CD groups not statistically significant (Figure 6I-j). Protein levels of both CTRP3 and SIRT1 in terminal ileal mucosa were likewise suppressed in both inactive and active CD groups relative to the control group, again with no statistically significant difference found between the active and inactive groups (Figure 6K-L). Finally, NF-κB p65 from whole-cell extracts of terminal ileal mucosa had similar expression levels across all three groups; however, phosphorylated NF-κB p65 was significantly higher in the CD groups relative to the control group (Figure 6K-L). Again, no statistically significant difference was observed in the phosphorylated NF-κB p65 between inactive and active CD groups; however, the ratios of phosphorylated NF-κB p65 to NF-κB p65 did reflect statistically significant differences, with the highest ratio in the active CD group, followed by the inactive CD group, and then the control group (Figure 6K-L). These data indicate that, as in DSS-induced mouse colitis models, CTRP3 attenuates intestinal inflammation in CD through SIRT1, which suppresses the pro-inflammatory transcriptional activity of phosphorylated NF-κB p65.

**Figure 6.**
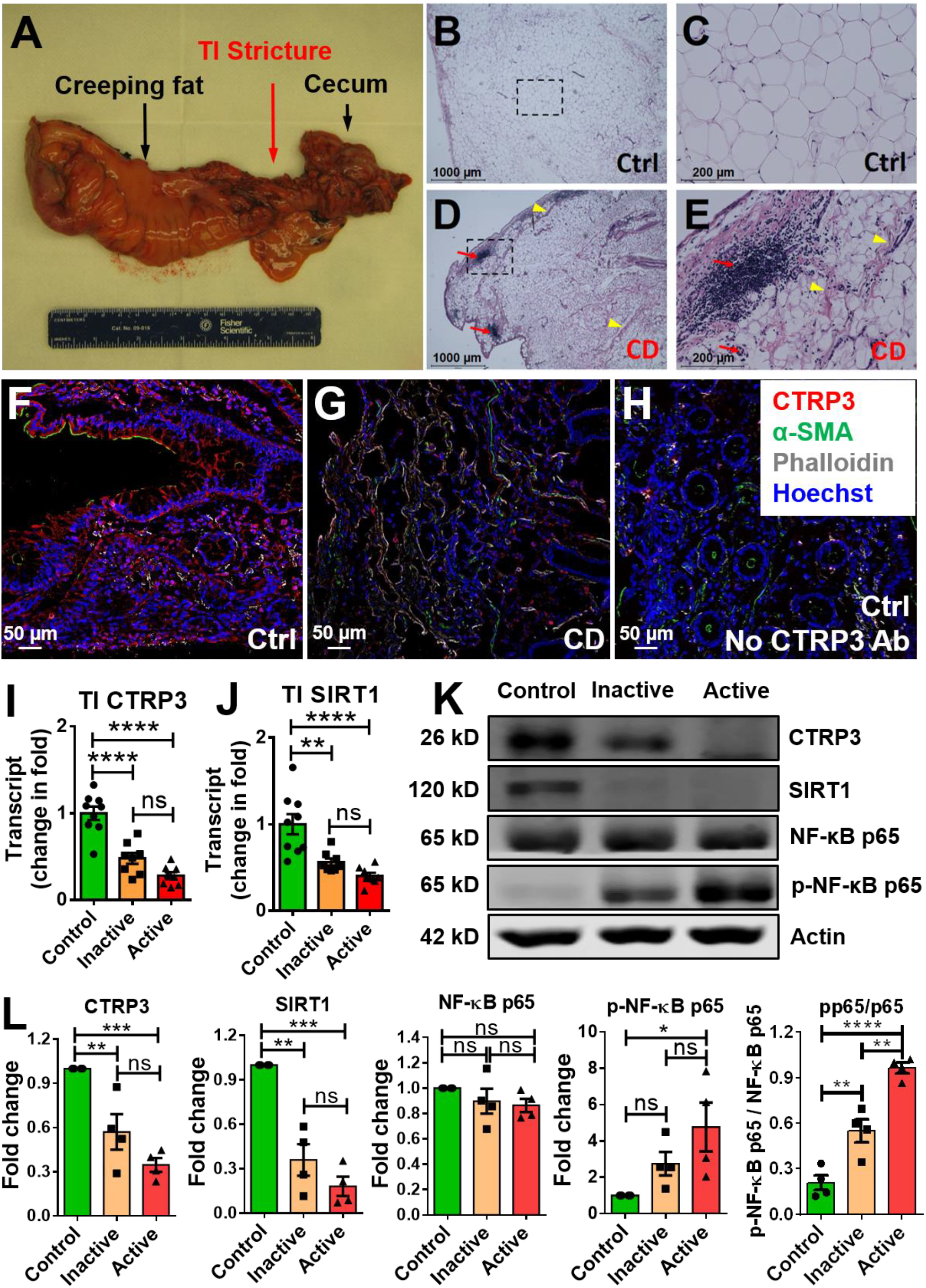
CTRP3 and SIRT1 are reduced in the terminal ileal mucosa of patients with active and inactive CD, resulting in increased phosphorylated NF-κB p65 transcriptional activity. **(A)** Macroscopic changes of creeping fat in CD. Surgical specimen from a CD patient with ileal stricture showing creeping fat wrapping around the inflamed terminal ileum (TI). **(B-E)** Histological changes of creeping fat in CD. H&E staining of mesenteric fat from CD patients (D, E) revealed smaller and more abundant adipocytes relative to control patients (B, C). Inflammatory infiltrates (red arrows); collagen deposition (yellow arrowheads). C/E are enlarged images of B/D boxed areas. **(F-H)** Immunofluorescence staining of human terminal ileal mucosa showed that CTRP3 protein was reduced in the active CD (G) compared with normal mucosa (F). H served as a negative control. α-SMA (smooth muscle actin), Phalloidin (F-actin) and Hoechst (nuclei). **(I-J)** mRNA levels of CTRP3 and SIRT1 were significantly lower in the terminal ileal mucosa from patients with inactive and active CD (n=8) compared to control patients (n =9). No statistically significant differences in the mRNA expression levels of both genes were observed between the inactive and active CD groups. Inactive and active epithelial tissue was obtained from the same 8 patients. Inactive/active groups normalized to the control group. **(K-L)** Protein levels of CTRP3 and SIRT1 were significantly lower in the terminal ileal mucosa from patients with inactive and active CD compared to control patients. The ratio of phosphorylated NF-κB p65 to NF-κB p65 from whole-cell extracts was highest in the active CD group and higher in the inactive group relative to the control group. Protein bands were quantified using ImageJ and normalized to β-actin (L). Results are shown as Mean ± SEM. \**P* < .05; \*\**P* < .01; \*\*\**P* < .001; **** *P* < .0001; ns, not significant (two-way ANOVA).

## Discussion

We are the first to demonstrate that a novel adipokine, CTRP3, attenuates intestinal inflammation in acute colitis mouse models. We found that DSS-induced acute colitis was much more severe in CTRP3 KO mice than in their WT littermates; conversely, colitis was much less severe in CTRP3 overexpressing Tg mice than in their WT littermates, suggesting that CTRP3 is a protective adipokine that suppresses intestinal inflammation under pathological conditions *in vivo*. Our study of human intestinal epithelial tissue yielded consistent results. We found that CD patients expressed lower epithelial levels of CTRP3 and SIRT1 than control patients, resulting in up-regulated pro-inflammatory NF-κB transcriptional activity.

Also significant is our finding that CTRP3 exerts an anti-inflammatory effect even under basal conditions. Although the clinical phenotypes and intestinal histologic scores of water-treated CTRP3 KO/Tg mice and water-treated WT littermates did not show significant differences (Figure 2B-D, 2F-G, 3B-D, 3F-G, and Supplementary Figure 3), CTRP3 levels correlated positively with colon length in water-treated CTRP3 KO and Tg mice. Further, in water-treated CTRP3 KO mice, colonic levels of SIRT1 were significantly reduced, while phosphorylated NF-κB p65, TNF-α, and IL-6 were significantly up-regulated, compared with their water-treated WT littermates (Figure 4B-F). This pattern was reversed, to a large extent, in water-treated CTRP3 overexpressing mice. Water-treated CTRP3 Tg mice expressed higher colonic levels of SIRT1 and lower levels of phosphorylated NF-κB p65 than their WT littermates (Figure 4B, E-E); levels of TNF-α, but not IL-6, were lower (Figure 5C-D). These findings suggest that CTRP3 not only attenuates pathological inflammation (DSS treatment in our colitis model), but also helps preserve an appropriate level of physiological intestinal inflammation (water treatment in our colitis model) —*i.e.,* the innate immune response of intestinal mucosa to non-pathological (or pre-pathological) aggravation.^61^ This is significant because IBD is increasingly thought to emanate from dysfunction in the innate immune system.^62–65^ It is likely that reduced CTRP3 levels in IBD patients are inadequate to enable the maintenance of homeostatic physiological inflammation. Future studies will investigate whether and how CTRP3 functions within the innate immune system (in both IBD and basal settings). More generally, these findings add to the increasing evidence that adipokines are critical to the maintenance of homeostasis in the intestinal epithelium.^66^

Our finding that DSS-treated mice experienced reductions in colonic CTRP3 levels raises important questions (Figure 4 and 5). It is possible that this was caused by pathophysiological changes echoing the reduced CTRP3 levels found among IBD patients (Figure 6). We located CTRP3 expression in intestinal epithelial cells, mesenteric adipocytes, and smooth muscle cells (Figure 1). In both our colitis mouse model and CD patient groups, intestinal epithelial cells and mesenteric adipocytes exhibited structural and functional changes (Supplementary Figure 2, Figure 6B-H). Smooth muscle cells were not examined in this study but are known to undergo morpho-functional changes in patients with CD, which often leads to intestinal fibrosis.67, 68 It is possible that intestinal inflammation results in structural and functional changes to these three local sources of CTRP3, which in turn further reduces CTRP3 production (as a negative feedback loop). 67 Of possible relevance, we found no statistically significant difference between the reduced CTRP3 levels in active and inactive CD patients (Figure 6I, K-L). This congruence could be viewed as evidence that IBD-related pathophysiological changes, rather than transient inflammatory activities, are etiologic to these reductions in CTRP3 levels. Future studies will probe this matter, which may implicate the important question of why CTRP3 levels are reduced among IBD patients.

Finally, our study helps to elucidate the molecular mechanisms through which CTRP3 attenuates intestinal inflammation by correlating concurrent changes in CTRP3 and SIRT1 levels with changes in levels of phosphorylated NF-κB p65 and, ultimately, intestinal inflammation. We are the first to show that CTRP3 exerts its anti-inflammatory effect through SIRT1/NF-κB signaling in intestinal inflammation, as has been observed in settings of acute pancreatitis and cardiac inflammation.^47–49^

Epithelial NF-κB activation and dysregulated cytokine production are well recognized in the pathogenesis of IBD.^52–54,69, 70^ In its quiescent state, NF-κB exists as an inactive cytoplasmic complex comprised of five transcription factor subunits.^70, 71^ Following stimulation, individual NF-κB subunits undergo phosphorylation, which optimizes their transactivation potential on a gene-specific basis. The phosphorylated subunits are translocated as dimers to the nucleus, where they effectuate gene expression.^70–72^ In IBD patients, phosphorylation of the NF-κB p65 subunit (which is translocated to the nucleus as a p65-p50 heterodimer) has been shown to up-regulate the expression of the pro-inflammatory cytokines TNF-α and IL-6.^53, 54^ SIRT1, a widely-expressed histone deacetylase, is known to mediate this process in a number of ways, including by influencing the nuclear translocation of phosphorylated NF-κB dimers, deacetylating them, and activating NF-κB inhibitors.^73–75^ Although not previously linked to CTRP3 in an intestinal inflammation setting, SIRT1 has been shown to attenuate intestinal inflammation in IBD by suppressing the transcriptional activity of phosphorylated NF-κB p65.^55–57^

In our studies of murine colitis models and patients with IBD, we demonstrated that CTRP3 regulates intestinal inflammation via SIRT1/NF-κB signaling. Consistent with previous studies^55–57^, we found that SIRT1 was reduced in the colonic tissue of DSS-treated mice and the terminal ileal mucosa of CD patients (both active and inactive). Our study links these reductions in SIRT1 levels to CTRP3. In both DSS- and water-treated groups, CTRP3 KO mice had significantly reduced colonic SIRT1 levels relative to their WT littermates (Figure 4B, E-F), while colonic SIRT1 levels were significantly increased in water-treated CTRP3 Tg mice relative to their WT littermates (Figure 5B, E-F). Likewise, reduced ileal CTRP3 levels in CD patients correlated with reduced SIRT1 levels (Figure 6I-L). Our study correlates these concurrent reductions in CTRP3 and SIRT1 levels with up-regulated intestinal levels of phosphorylated NF-κB p65 (Figure 4E-F, 6I-L) and the expression of pro-inflammatory cytokines TNF-α and IL-6 (Figure 4C-D). Although the precise signaling relationship between CTRP3 and SIRT1 remains unknown, our findings indicate that CTRP3 suppresses intestinal inflammation through the SIRT1/NF-κB signaling axis — including through SIRT1-mediated, NF-κB p65 transcriptional activity (Figure 7).

**Figure 7.**
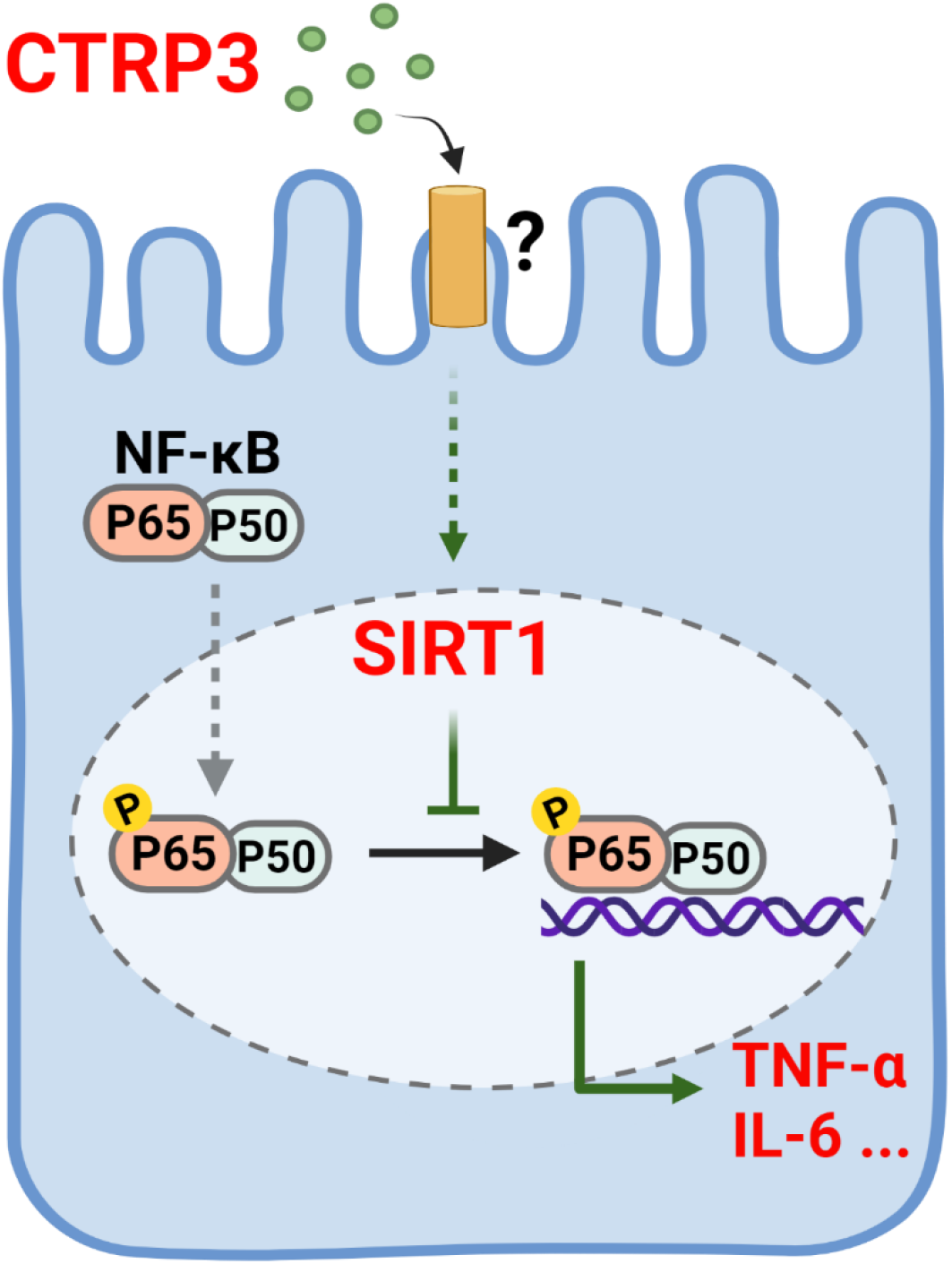
Proposed model of CTRP3-mediated anti-inflammatory signaling in intestinal epithelial cells. Through an unknown receptor, CTRP3 (green dots) attenuates intestinal epithelial inflammation through SIRT1, which suppresses transcriptional activity of phosphorylated NF-κB p65, resulting in down-regulated expression of pro-inflammatory cytokines TNF-α and IL-6.

In conclusion, our findings indicate that CTRP3 attenuates intestinal inflammation in mouse colitis models and CD patients through SIRT1/NF-κB signaling. Future studies are needed to fully dissect the mechanisms through which CTRP3 activates this signaling, the specific tissue sources of CTRP3 that influence intestinal inflammation, and whether the administration of recombinant CTRP3, SIRT1 agonists, and/or NF-κB antagonists have translational potential as IBD therapies.

## Abbreviations used in this paper

CD: Crohn’s disease
CTRP3: C1q/tumor necrosis factor-related protein 3
DSS: dextran sulfate sodium
IL-6: interleukin-6
IBD: inflammatory bowel disease
NF-κB: nuclear factor kappa-light-chain-enhancer of activated B cells
SIRT1: Sirtuin 1
TNF-α: tumor necrosis factor-α
smFISH: single-molecule RNA fluorescence *in situ* hybridization
UC: ulcerative colitis

## Acknowledgments

We thank Chung-Ming Tse, Michael Goggins, Stephen Meltzer, Tatianna Larman, James Hamilton, James Potter, and Varsha Singh for their advice and comments on the manuscript. We thank Mark Lazarev, Bashar Safar, and Chady Atallah for their assistance in obtaining human tissue samples. We thank Liping Li, George McNamara, Ruxian Lin, and Denise Chesner for their technical assistance.

The current address for Gangping Li is: Division of Gastroenterology, Union Hospital, Tongji Medical College, Huazhong University of Science and Technology, 1277 Jiefang Avenue, Wuhan, China.

The current address for Lingling Xian is: Department of Pathology, University of South Alabama, 2451 University Hospital Drive, Mobile, AL

## CRediT Authorship contributions

➢ Huimin Yu, MD, PhD (Conceptualization: Lead; Data curation: Lead; Formal analysis: Lead; Funding acquisition: Lead; Investigation: Lead; Methodology: Lead; Project administration: Lead; Resources: Lead; Supervision: Lead; Validation: Lead; Visualization: Lead; Software: Supporting; Writing - original draft: Lead; Writing – review & editing: Lead)
➢ Zixin Zhang, MS (Data curation: Lead; Formal analysis: Lead; Investigation: Lead; Visualization: Lead; Software: Lead; Writing – original draft: Supporting)
➢ Gangping Li, MD, PhD (Data curation: Lead; Formal analysis: Lead; Investigation: Lead; Visualization: Supporting; Software: Lead)
➢ Yan Feng, MD, PhD (Conceptualization: Supporting; Data curation: Supporting; Formal analysis: Supporting; Investigation: Supporting; Methodology: Supporting; Visualization: Supporting; Software: Supporting; Writing – review & editing: Supporting)
➢ Lingling Xian, MD, PhD (Conceptualization: Supporting; Data curation: Supporting; Formal analysis: Supporting; Investigation: Supporting; Methodology: Supporting; Visualization: Supporting; Software: Supporting; Writing – review & editing: Supporting)
➢ Fatemeh J Bakhsh, PhD (Investigation: Supporting)
➢ Dongqing Xu, PhD (Investigation: Supporting)
➢ Cheng Xu, PhD (Resources: Supporting)
➢ Tyrus Vong, BS (Resources: Supporting; Data curation: Supporting)
➢ Bin Wu, PhD (Methodology: Supporting)
➢ Florin M Selaru, MD, MBA (Resources: Supporting; Writing – review & editing: Supporting)
➢ Fengyi Wan, PhD (Formal analysis: Supporting; Methodology: Supporting; Resources: Supporting; Writing - review & editing: Supporting)
➢ G. William Wong, PhD (Conceptualization: Supporting; Funding acquisition: Supporting; Formal analysis: Supporting; Methodology: Supporting; Resources: Supporting; Writing - review & editing: Supporting)
➢ Mark Donowitz, MD (Formal analysis: Supporting; Methodology: Supporting; Resources: Supporting; Supervision: Supporting; Writing – review & editing: Supporting)

## Supplemental Figures

**Supplemental Figure 1.**
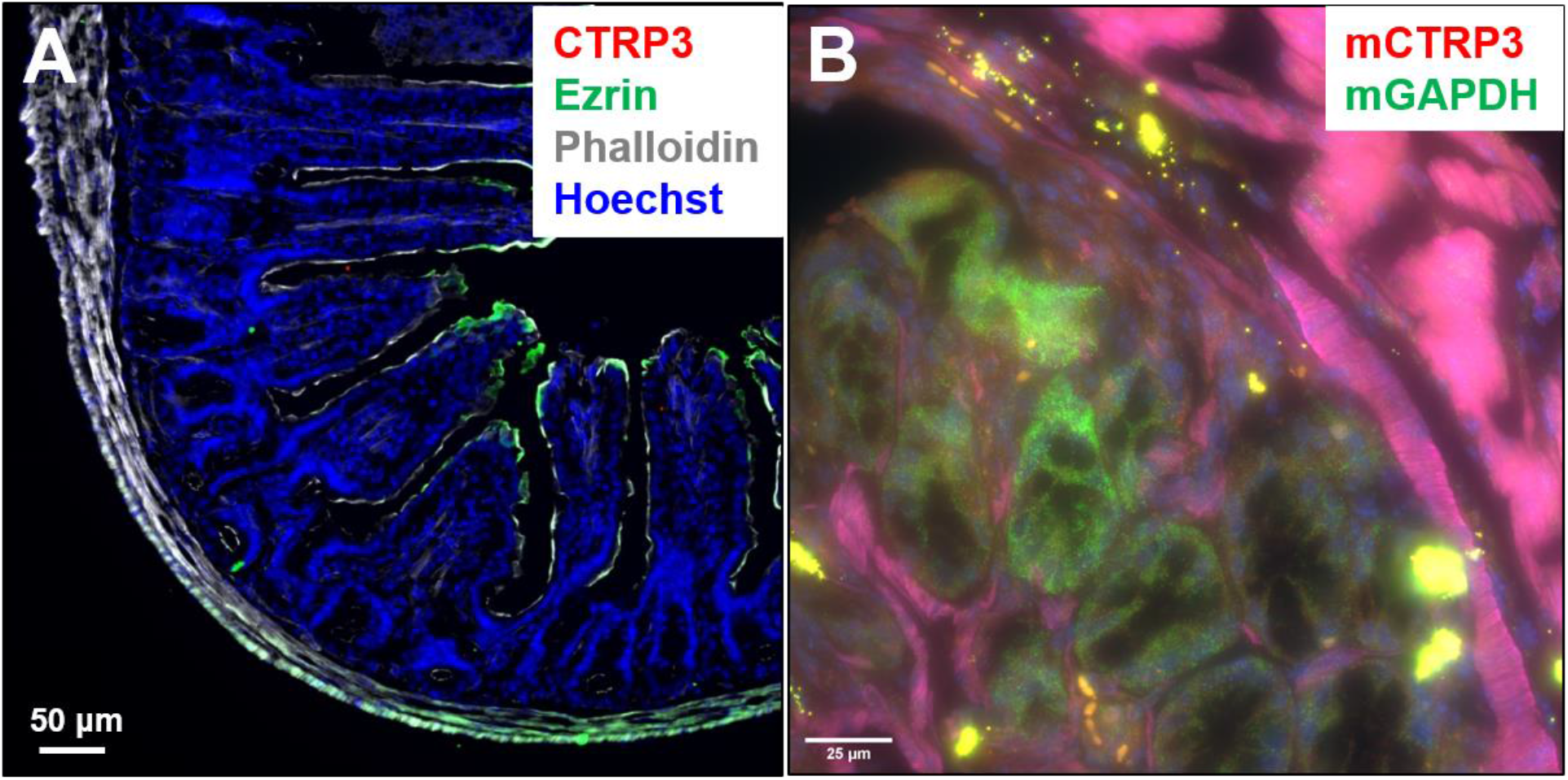
Colonic tissue from CTRP3 KO mice was used as negative controls in **(A)** immunofluorescence staining and **(B)** single-molecule RNA fluorescence *in situ* hybridization (smFISH). mGAPDH (green dots, a positive control for smFISH).

**Supplemental Figure 2.**
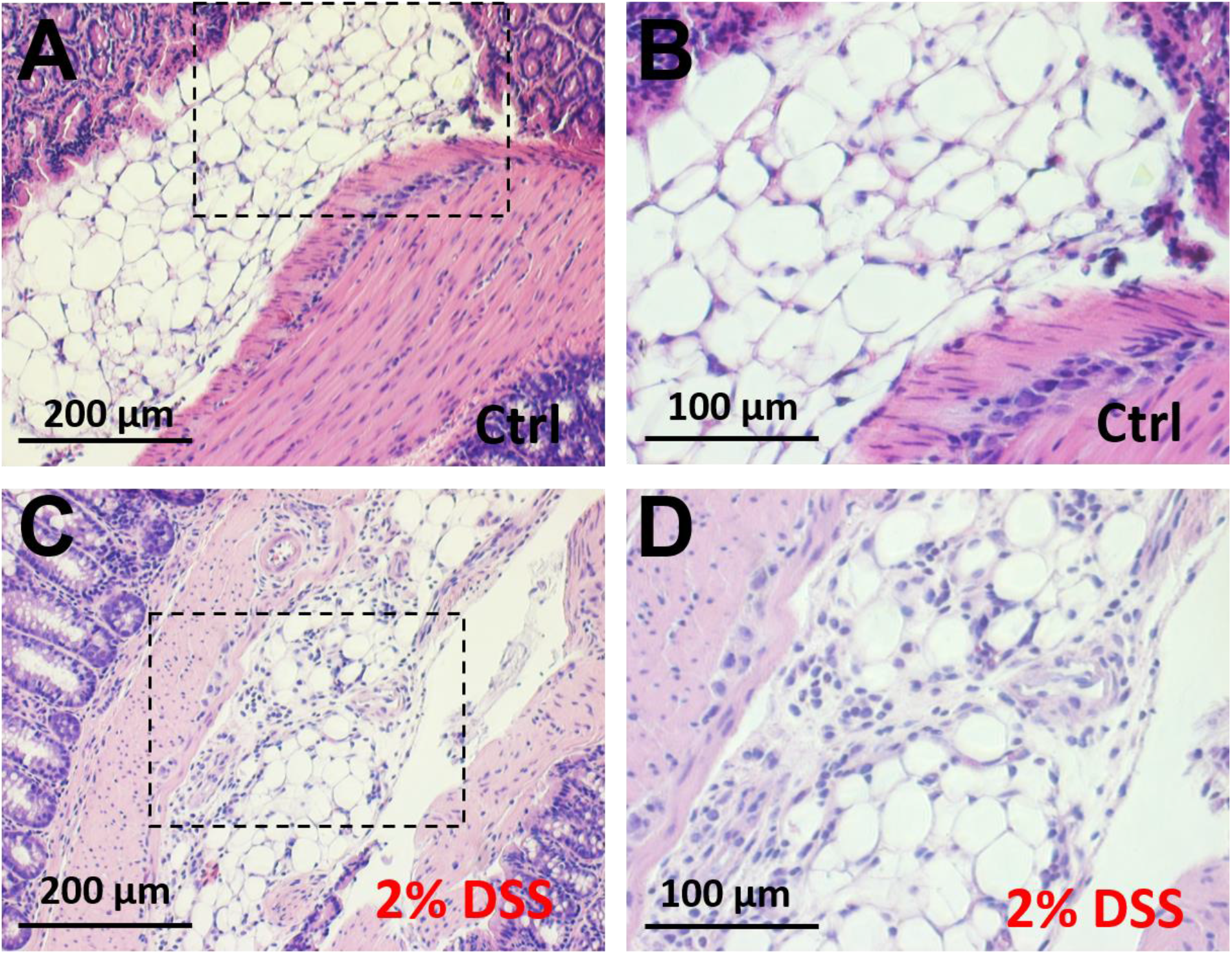
Mesenteric fat exhibits adipocyte hyperplasia in acute DSS-induced colitis. H&E staining of mesenteric fat collected from WT mice treated with water **(A, B)** or 2% DSS for only 6 days **(C, D)** revealed that DSS-treated mice had smaller and more abundant adipocytes and increased inflammatory cell infiltrates. B/D are enlarged images of A/C boxed areas.

**Supplemental Figure 3.**
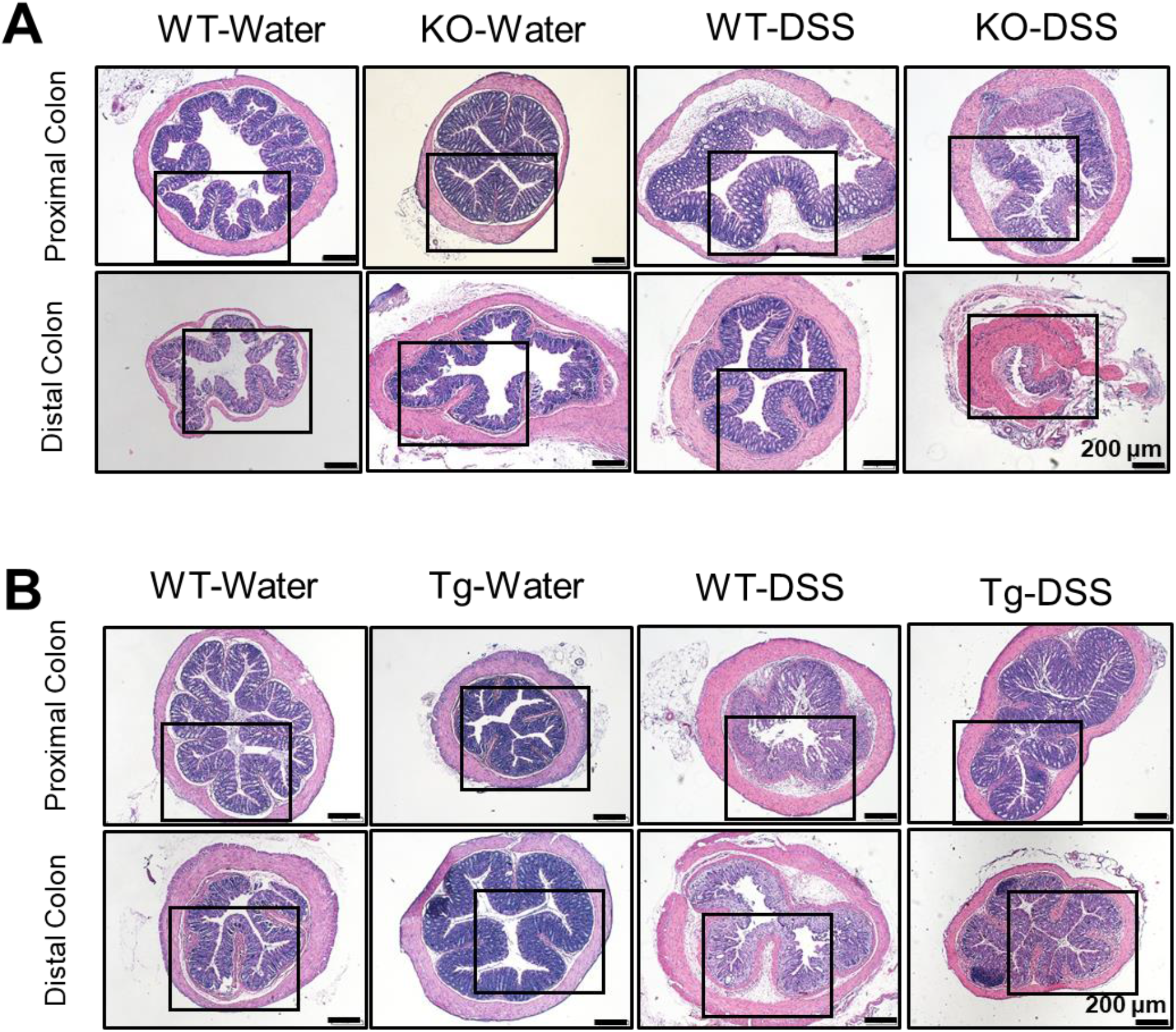
Representative images of H&E staining of the proximal and distal colon of water- and DSS-treated **CTRP3 KO (A)** and **CTRP3 Tg (B)** mice. N = 5 −10 mice/group. **(A)** DSS-treated CTRP3 KO mice exhibited significantly more tissue damage and inflammatory cell infiltrates than in DSS-treated WT mice. Water-treated CTRP3 KO and WT mice showed no difference in histologic analysis. **(B)** DSS-treated CTRP3 Tg mice exhibited significantly less tissue damage and fewer inflammatory cell infiltrates than their WT littermates. Water-treated CTRP3 Tg and WT mice showed no difference in histologic analysis.

**Supplemental Figure 4.**
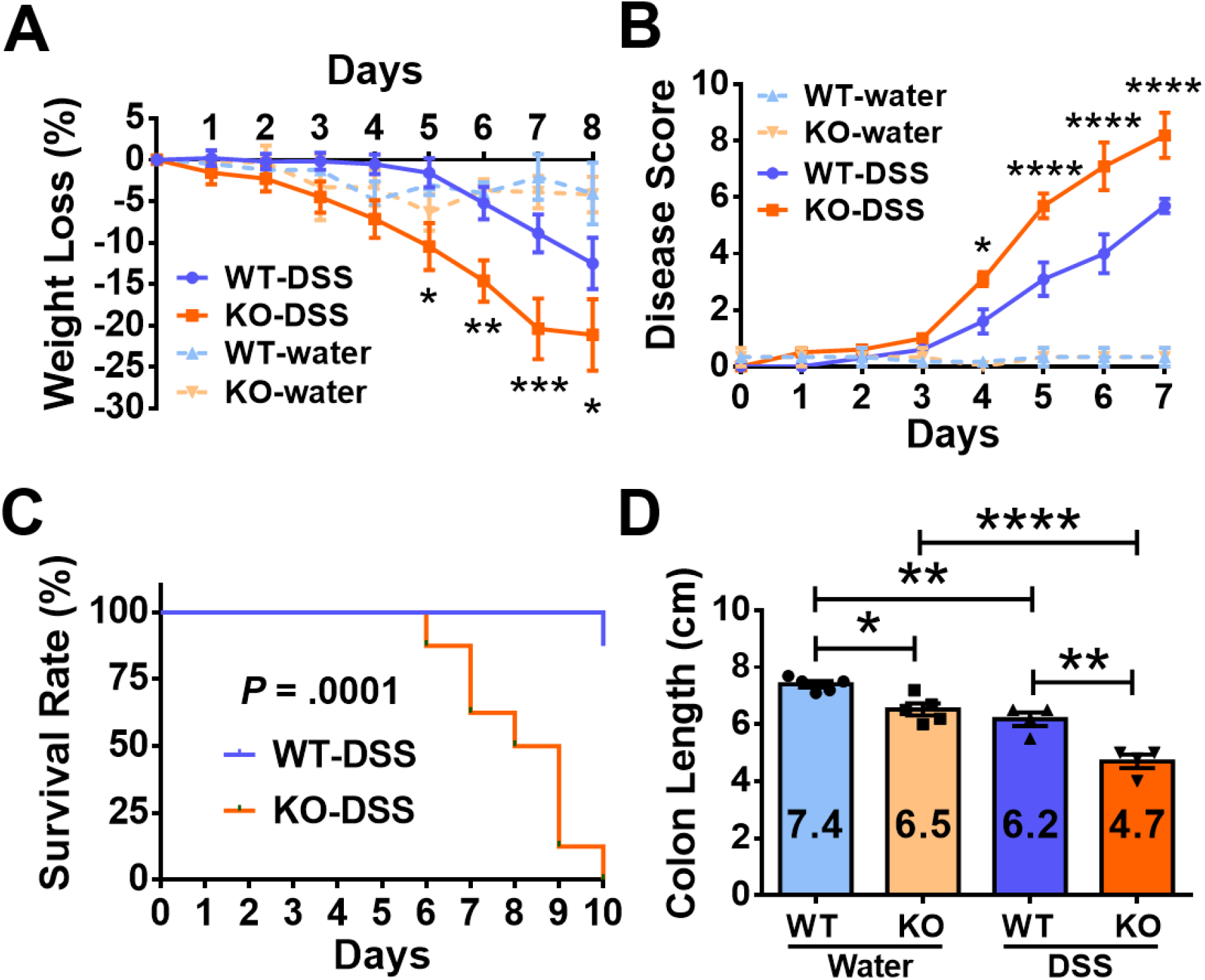
Female CTRP3 KO mice develop more severe acute colitis with 1.2% DSS than WT littermates in acute DSS-induced colitis models. **(A-B)** DSS-treated CTRP3 KO mice had more pronounced weight loss and developed more severe colitis relative to DSS-treated WT littermates. Water-treated CTRP3 KO and WT mice showed no difference in weight loss and clinical colitis phenotypes. (N = 5 in each DSS-treated group and N = 3 in each water-treated group). Statistical comparison was between DSS-treated CTRP3 KO and WT mice. **(C)** DSS-treated CTRP3 KO mice displayed a lower survival rate relative to DSS-treated WT littermates. Log-rank Mantel-Cox test was used for survival analysis. (N = 8 in each group) **(D)** The colons of CTRP3 KO mice were significantly shorter than those of WT littermates in both water- and DSS-treated groups. (N = 4 in each DSS-treated group and N = 5 in each water-treated group). Results are shown as means ± SEM. \**P* <.05; \*\**P* < .01; \*\*\**P* < .001; **** *P* < .0001 (two-way ANOVA except for the survival curve).

**Supplemental Figure 5.**
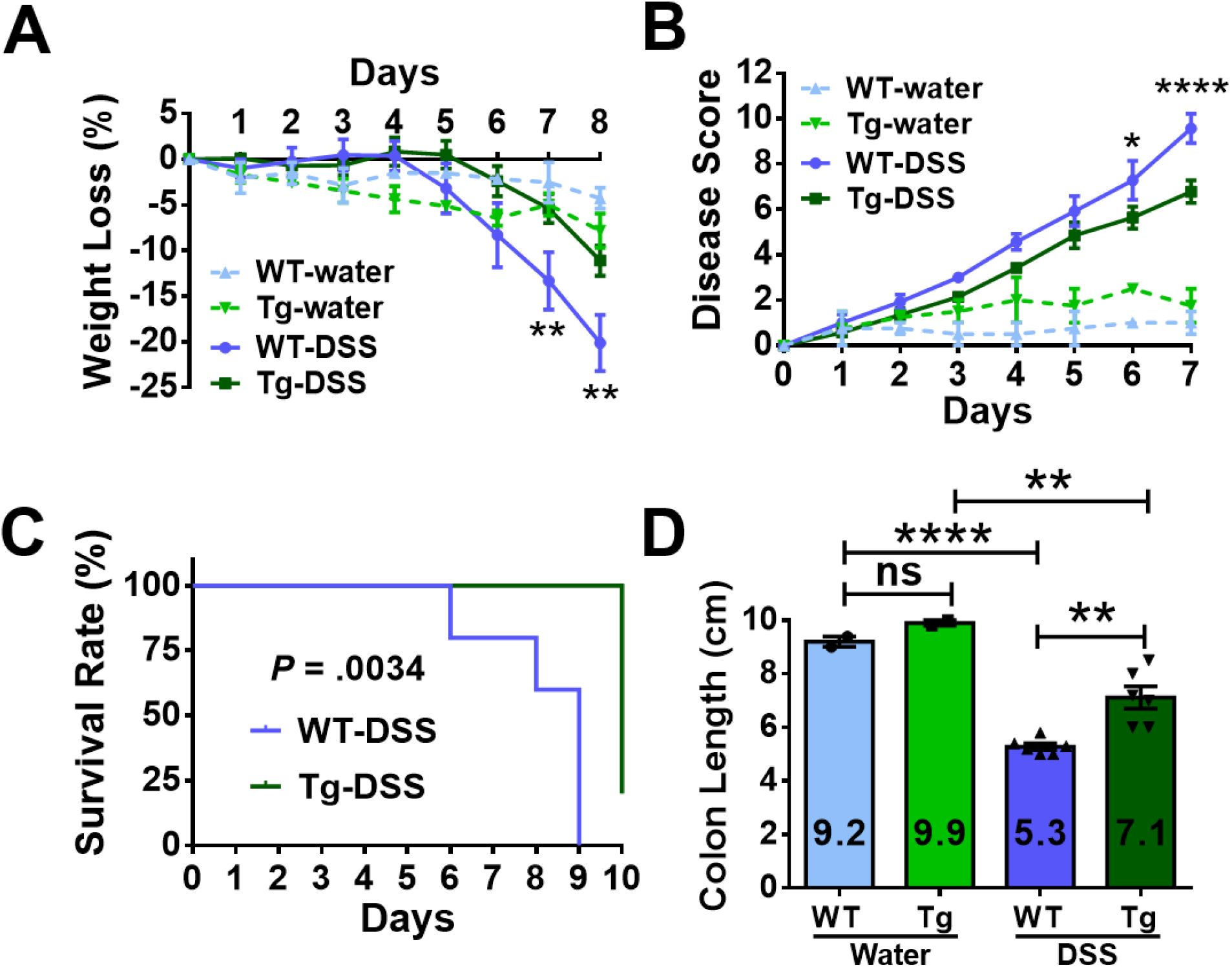
Female CTRP3 Tg mice develop less severe colitis with 2% DSS relative to DSS-treated WT littermates in acute DSS-induced colitis models. **(A-B)** DSS-treated CTRP3 Tg mice had less pronounced weight loss and developed less severe colitis relative to DSS-treated WT littermates. Water-treated CTRP3 Tg and WT mice showed no difference in weight loss and clinical colitis phenotypes. (N = 6 in each DSS-treated group and N = 2 in each water-treated group). Statistical comparison was between DSS-treated CTRP3 KO and WT mice. **(C)** DSS-treated CTRP3 Tg mice displayed a higher survival rate relative to DSS-treated WT littermates. Log-rank Mantel-Cox test was used for survival analysis. (N = 5 in each group) **(D)** The colons of CTRP3 Tg mice were significantly longer than those of WT littermates in both water- and DSS-treated groups. (N = 6 in each DSS-treated group and N = 2 in each water-treated group). Results are shown as means ± SEM. \**P* <.05; \*\**P* <0 .01; \*\*\**P* < .001; **** *P* < .0001 (two-way ANOVA except for the survival curve).

**Supplemental Figure 6.**
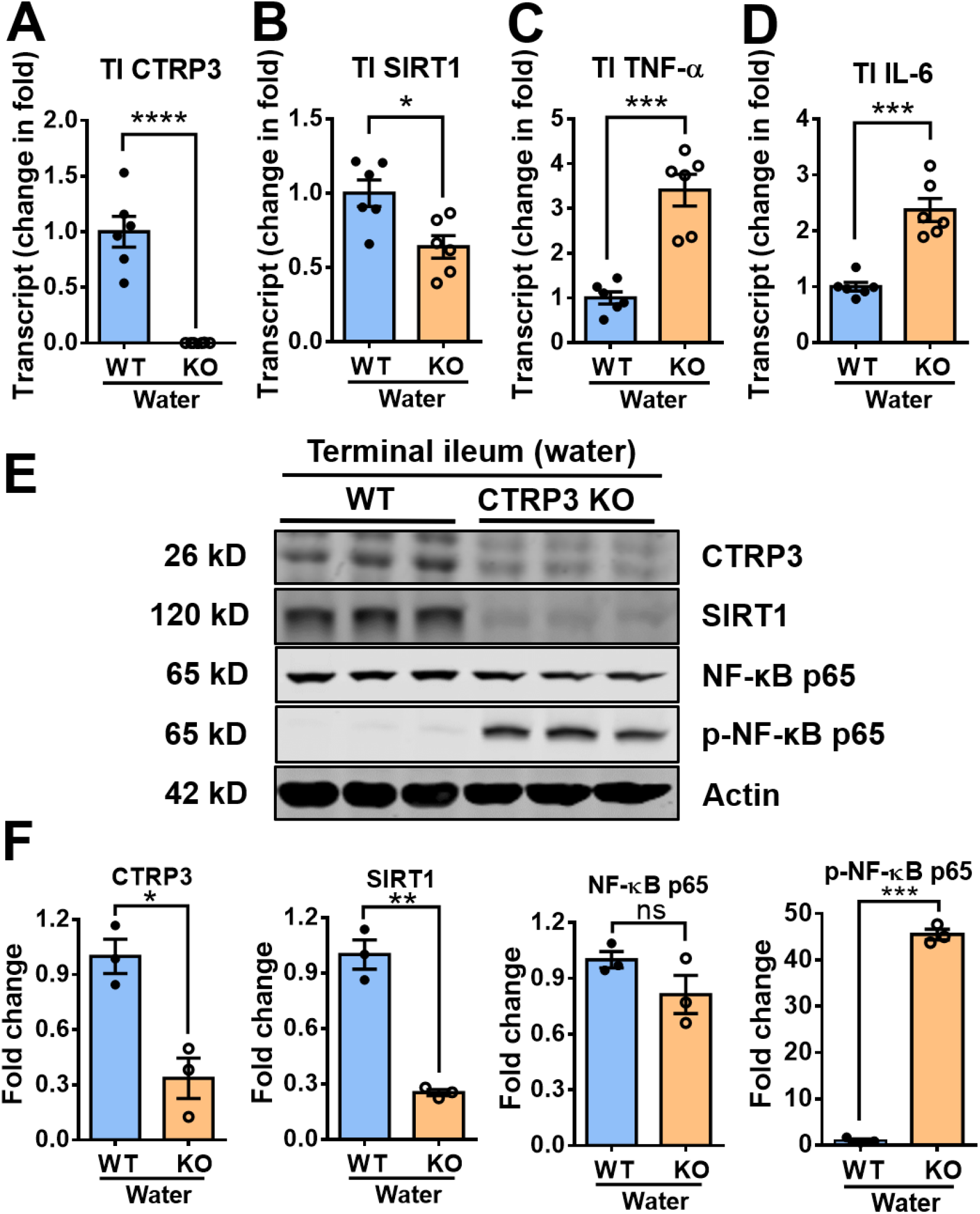
Terminal ileal tissue of CTRP3 KO mice exhibits decreased SIRT1 and increased phosphorylated NF-κB p65 transcriptional activity (water-treated). **(A-B)** mRNA levels of CTRP3 and SIRT1 were significantly lower in CTRP3 KO mice relative to WT mice. **(C-D)** TNF-α and IL-6 mRNA levels were up-regulated in CTRP3 KO mice relative to WT mice. **(E-F)** SIRT1 protein levels were lower in CTRP3 KO mice relative to WT mice. NF-κB p65 from whole-cell extracts had similar expression levels in the two groups; however, phosphorylated NF-kB p65 was up-regulated in the TI of CTRP3 KO mice. 3 mice/group shown in (E). Protein band were quantified using ImageJ and normalized to β-actin (F). Results are shown as means ± SEM; N = 6 mice/group. \**P* < .05; \*\**P* < .01; \*\*\**P* < .001; **** *P* < .0001; ns, not significant (Student’s t-test).

## Supplementary Materials and Methods

**Supplementary Table 1.** Criteria for the clinical disease score

|  | Weight loss | Stool consistency | Rectal bleeding |
| --- | --- | --- | --- |
| 0 | None | Normal | No bleeding |
| 1 | 1% - 5% weight loss | Slight diarrhea | POBT detected |
| 2 | 5% - 10% weight loss | Diarrhea | Slight bleeding |
| 3 | 10% - 15% weight loss | Slight watery diarrhea | Bleeding |
| 4 | 15% - 20% weight loss | Watery diarrhea | Gross bleeding |

**Supplementary Table 2.** Criteria for the histological score

|  | Tissue damage | Inflammatory cell infiltration |
| --- | --- | --- |
| 0 | < 5 % | Infrequent |
| 1 | 5 – 25 % | Increased, some neutrophils |
| 2 | 25 – 50 % | Submucosal, inflammatory cell clusters |
| 3 | 50 – 75 % | Infiltrations muscular |
| 4 | > 75 % | Transmural |

**Supplementary Table 3.**
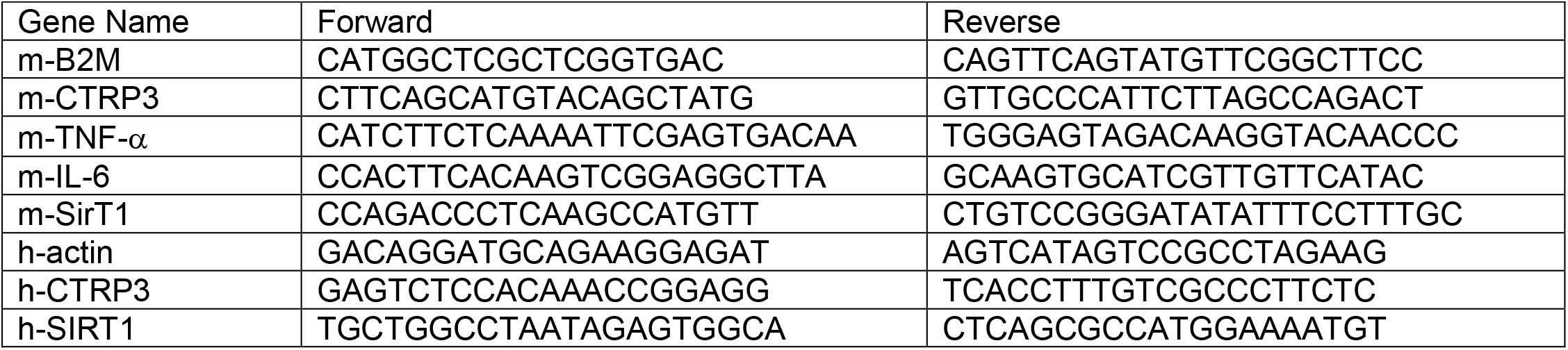
Primers information for RT-qPCR

### Mouse genotyping

Mouse tail DNA for genotyping was isolated via phenol/chloroform extraction and isopropanol precipitation according to a standard protocol. Standard polymerase chain reactions (PCRs) were performed with Hot Start Go Taq polymerase (Promega, Madison, WI) by using the following primers in Supplementary Table 4.

**Supplementary Table 4.**
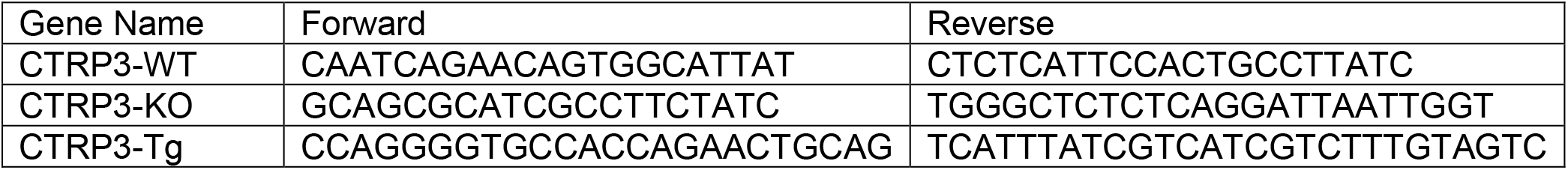
Primers information for genotyping

